# Synthetic Data Resource and Benchmarks for Time Cell Analysis and Detection Algorithms

**DOI:** 10.1101/2022.01.01.474717

**Authors:** Kambadur Gundu Ananthamurthy, Upinder S Bhalla

## Abstract

Hippocampal CA1 cells take part in reliable, time-locked activity sequences in tasks that involve an association between temporally separated stimuli, in a manner that tiles the interval between the stimuli. Such cells have been termed time cells. Here we adopt a first-principles approach to comparing diverse analysis and detection algorithms for identifying time cells. We generated synthetic activity datasets using calcium signals recorded *in vivo* from the mouse hippocampus using 2-Photon imaging, as template response waveforms. We assigned known, ground truth values to perturbations applied to perfect activity signals, including noise, calcium event width, timing imprecision, hit-trial ratio and background (untuned) activity. We tested a range of published and new algorithms and their variants on this dataset. We find that most algorithms correctly classify over 80% of cells, but have different balances between true and false positives, and different sensitivity to the five categories of perturbation. Reassuringly, most methods are reasonably robust to perturbations, including background activity, and show good concordance in classification of time cells. The same algorithms were also used to analyse and identify time cells in experimental physiology datasets recorded *in vivo* and most show good concordance.

**Significance Statement:** Numerous approaches have been developed to analyze time cells and neuronal activity sequences, but it is not clear if their classifications match, nor how sensitive they are to various sources of data variability. We provide two main contributions to address this: 1) A resource to generate ground truth labelled synthetic 2-P Calcium activity data with defined distributions for confounds such as noise and background activity, and 2) a survey of several methods for analyzing time-cell data using our synthetic data as ground truth. As a further resource, we provide a library of efficient C++ implementations of several algorithms with a Python interface. The synthetic dataset and its generation code are useful for profiling future methods, testing analysis toolchains, and as input to computational and experimental models of sequence detection.

## Introduction

The mammalian hippocampus is important for the formation of several kinds of memory, one of which is the association between stimuli occurring separately in time. Time cells were originally described using tuning curves from single-unit recordings of cellular activity when rats ran on a running wheel in between behavioural decisions (Pastalkova et al. 2008). These cells exhibited time tuning of the order of seconds. Several further studies have shown that small populations of hippocampal CA1 cells fire in time-locked sequences, “bridging” the time gap between stimulus and response in temporal delay tasks lasting several seconds (Kraus et al., 2013; MacDonald et al., 2011, 2013; Pastalkova et al., 2008). Cellular calcium imaging studies have also been used to report time cells, albeit at slower sampling rate (Mau et al., 2018; Modi et al., 2014). For example, similar interval tiling properties of hippocampal CA1 neurons were observed on much shorter, 500 ms time-scales in a Trace Eye-Blink Conditioning (TEC) task (Modi et al., 2014). Spontaneous sequential activity has also been reported in free-running animals (Villette et al., 2015)□. Such cells with a well-defined temporal firing field are commonly termed time cells (Eichenbaum, 2017; MacDonald et al., 2011)□. However, there is a wide diversity of methods used to detect and characterise time cells, and it is not clear how consistent these methods are in classifying cells as time cells. It is also unclear how sensitive each method may be to a range of physiological sources of variability and noise. A consistent set of benchmarks of classification performance is necessary to draw accurate and comparable conclusions from real physiology data across different methods and different laboratories. Our approach in the current study is not prescriptive, but pragmatic: we ask how existing methods work when we already know exactly which cells are time cells, and we determine how well each method deals with imperfect data.

The major approaches used to identifying time cells are tuning curves (peristimulus time histograms), temporal information, principal component analysis with time offset, support vector machines, and bootstrap analysis of activity peaks. Several studies have used a temporal delay task lasting several seconds, in which a rat runs on a treadmill during the delay period. A Temporal Information metric (Mau et al., 2018) has been used to find individual time cells in such tasks. A distinct task involves monitoring recurrent sequences of activity during free-running treadmill recordings. Such datasets have been analysed using Offset Principal Component Analysis (Kaifosh et al., 2013; Malvache et al., 2016; Villette et al., 2015), to first denoise 2-P data, establish correlation coefficients, and detect hippocampal CA1 sequences. Time cells have also been reported for much shorter duration tasks (~500 ms) such as hippocampus-dependent trace conditioning (Modi et al., 2014; Tseng et al., 2004). Time cells in these 2-P datasets were identified using yet another method, in which bootstrapping was used to determine if peak activity at a given time was different from chance. This method was termed Ratio of Ridge/Background (Modi et al., 2014). Yet other methods have utilised support vector machines to categorise time cells (Ahmed et al., 2020). Additionally, while the applicability of a variety of algorithms for place cell detection has been previously compared (Souza et al., 2018), we have focussed on methods which are fully automatable and which scale well to large datasets, specifically comparing algorithms to detect time cells.

Time cell detection is closely related to sequence detection, which has been fraught with statistical challenges. For example, detection of synfire chains has been the subject of some debate (Ikegaya et al., 2004; Lee & Wilson, 2004; Mokeichev et al., 2007; Schrader et al., 2008). Time cell detection is usually easier, in that in most experiments there is a well-defined initiating stimulus and a known delay or trace phase (however, see Villette et al., 2015). For any cell identified as a time cell, it is desirable to define a score to measure quality or reliability along with decodable time. Hence it is also valuable to be able to compare the score of a time cell across recordings and even between groups, using well defined, analog measures. Each algorithm currently used in the literature implements a different scoring method and it is as yet unclear if comparable results would be observed with other metrics.

In the current study we compare these diverse methods by estimating their performance on synthetic test datasets where we controlled all features of the data, including the identity and timing of each time cell. The development of a synthetic dataset serves two purposes. First, it facilitates principled comparison of different methods, since the ground truth is known. Second, it facilitates an analysis over many dimensions of input variance, corresponding to very different experimental and neural circuit contexts. Richness in variety of input data allows for better sampling of the performance of the analyses under many potential conditions. We have explored variance along the key dimensions of Noise, Timing Imprecision, Signal Widths, Frequency of Occurrence, as well as several others. To strengthen the applicability of this synthetic data resource to real data, our generated output uses sampled experimental data.

Our experimental data, synthetic dataset, and code base are intended to be a resource for algorithm testing and optimization.

## Methods

### Animals, Chronic Implants, and Behavioural Training

All animal procedures were performed in accordance with the National Centre for 114 Biological Sciences Institutional Animal Ethics Committee (Project ID: NCBS115 IAE-2016/20(M)), in accordance with the guidelines of the Government of India (Animal Facility CPCSEA registration number 109/1999/CPCSEA) and equivalent guidelines of the Society for Neuroscience.

To chronically monitor the activity of the same population of Hippocampal CA1 cells, we implanted 2-4 month old male and female GCaMP6f mice [Tg(Thy1-GCaMP6f)GP5.17Dkim JAX stock #025393] with an optical window and head-bar using a protocol adapted from previously published methods (Dombeck et al., 2010)□. Briefly, anaesthesia was induced with 2-3% Isoflurane in a chamber, and subsequently maintained (breathing rate of ~1 Hz) with 1-2% Isoflurane, directly to the mouse’s nose using an inverted pipette tip. Surgery was performed on a temperature-controlled table, maintained at 36.5 °C, while the anaesthetised animal was cheek-clamped. After a haircut, a ~5 cm piece of scalp was cut open to reveal the skull. A ~3mm circular craniotomy was then performed at a position 2 mm caudal and ~1.5 mm lateral to Bregma, on the left hemisphere. After gently tearing off the Dura, the underlying cortex was carefully aspirated till the Corpus Callosum (CC) layer, clearing out any blood using repeated washes of Cortex Buffer (Modi et al. 2014). A small thickness of corpus callosum fibres were then carefully aspirated till horizontal CC fibres were sparse but visible. The Cortex Buffer was then carefully suctioned out to dry the exposure till tacky. The exposure was then quickly sealed using a thin layer of Kwik-Sil and a coverslip attached to the bottom of a 3mm steel cannula. This preparation left the CA1 cell body layer ~200 μm below the most exposed tissue. Finally, an imaging head-bar was surgically implanted and fixed to the scalp, using dental cement and skull screws, before the animal was brought out of anaesthesia.

The animals were allowed to recover for 1-5 days after implantation, with a further 3-4 day habituation to the rig. Following this simultaneous behavioural training and 2-P *in vivo* imaging was conducted.

### Trace Eyeblink Conditioning (TEC)

We standardized a multi-session Trace Eyeblink Conditioning (TEC) paradigm to train head-fixed mice, based on previous literature □(Siegel et al., 2015). TEC involves an association between a previously neutral Conditioned Stimulus (CS) with an eye-blink inducing Unconditioned Stimulus (US), across an intervening, stimulus-free, Trace Interval. Training involved 60 trials per session, one session a day, for ~2 weeks. The CS was a 50 ms blue LED flash while the US was a 50 ms air-puff to the left eye. The stimulus free Trace Interval was 250-750 ms long, depending on the session. Additionally, a pseudorandom 10% of the trials were CS-only Probe Trials (no US) to test for learning. All behaviour routines were controlled by programmed Arduinos. Eye-blinks were measured for every trial, by video camera (Point Grey Chameleon3 1.3 MP Monochrome USB3.0) based detection.

The Conditioned Response (CR) is observed as a pre-emptive blink before the US is delivered, in animals that learn the task. The analysis of the behavioural data was performed using custom written MATLAB scripts. In brief, each frame for every trial was,

1. Cropped to get the eye,
2. Binarized to get the pixels defining just the eye, and finally
3. Given an FEC score from 0 to 1 (see below).

Every trial was then scored as a hit or miss, using the result of a 2 sample KS-Test between the FEC during the Trace and Pre-CS period (1% significance). The performance of an animal for a session was then established as the percentage of Hit Trials/Total Trials. Definitions –

FEC: The Fraction of Eye-Closed is estimated by counting the pixels defining the eye in every image of a time series, normalized by the maximum number of pixels defining the eye, in that session. Thus, every frame was given an analog score from 0 to 1, where,

- 0: fully opened eye
- 1: fully closed eye.

CR: The Conditioned Response is the eye-closing transition during the Trace period UR: The Unconditioned Response is the eye-closing transition when the US is delivered

Performance: Percentage of Hit Trials/Total Trials This allowed us to observe how the animals perform during and across sessions.

### Two-Photon Imaging

We used a custom-built two photon laser-scanning microscope (Modi et al., 2014) to record calcium activity from 100-150 Hippocampal CA1 cell bodies in vivo, at cellular resolution. We performed galvo-scans through the imaging window, over a field of view of ~100×100 μm^2^, at 14.5 Hz, during TEC (Figure 1A). An Arduino microcontroller was used to control the behaviour routines, and it additionally sent a TTL trigger to initiate the imaging trials. The behaviour and imaging were conducted simultaneously to record calcium activity when the animal was learning the task

**Figure 1:**
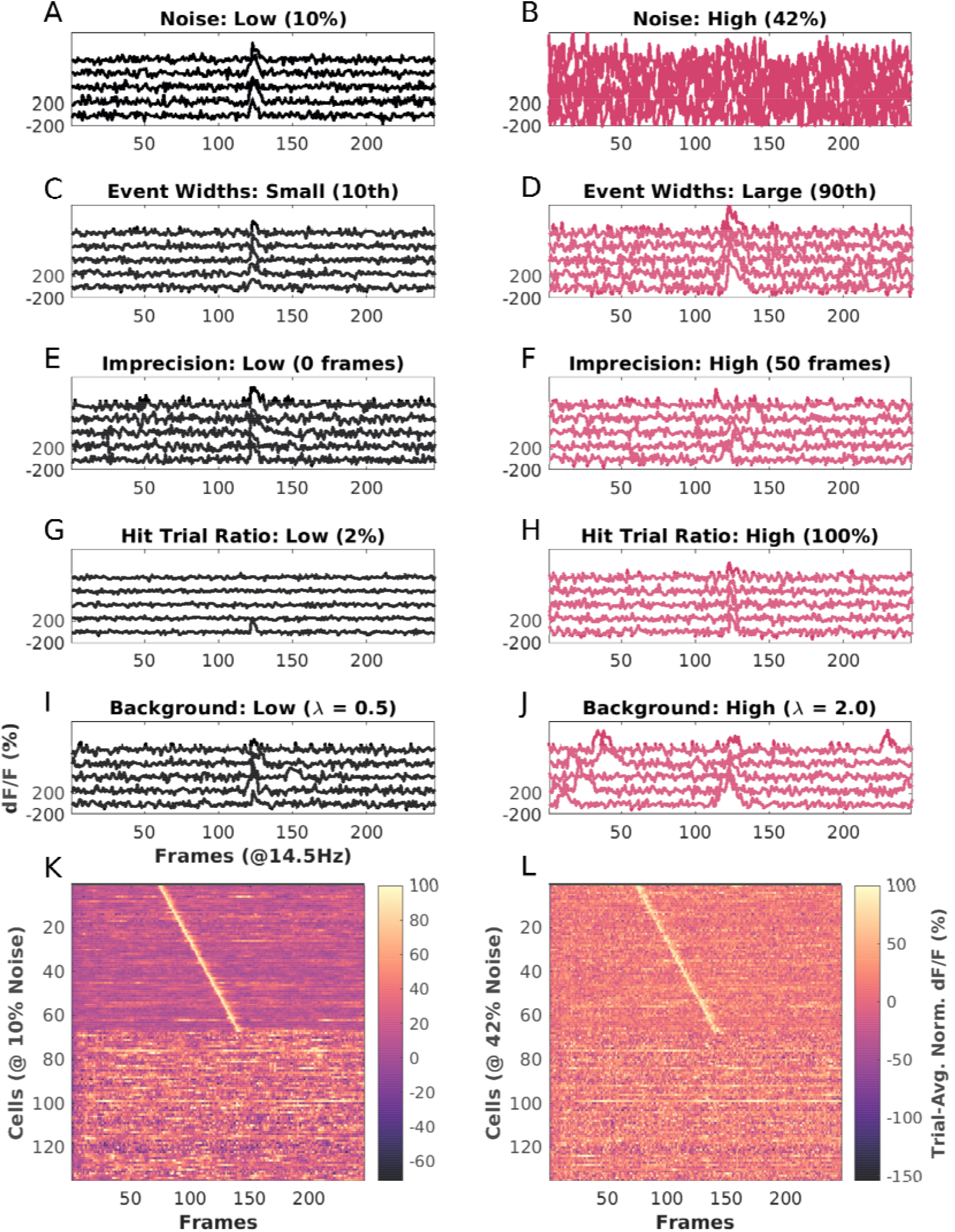
Key features of synthetic datasets. Left, black panels: low range of features. Right, red panels: High range of features. A: Noise = 10% B: Noise = 42%, C: Event Width: 10th percentile +/− 1 std. dev., D: Event Width 90th percentile +/ 1 std. dev. E: Imprecision, 0 frames FWHM F: Imprecision: Right: 50 frames FWHM, G: Hit Trial Ratio from 0 to 2%. H: Hit Trial ratio from 0 to 100%. I-J: Background Activity with the number of background spikes per background sampled from a Poisson Distribution for with mean (λ), for I: λ=0.5 (Low), and J: λ=3.0 (High). K-L: Trial-Averaged. Calcium Traces from example synthetic datasets of 135 neurons, displayed as heatmap sorted by time of peak Ca signal, I: Baseline Physiology synthetic data trial-average with 10% Noise (Low) and low background activity (λ=0.5 events/trial), and L: Same as K with 42% Noise (High) and higher background activity (λ=3.0 events/trial). In both cases 50% of the cells (top 67) are time cells and the remainder are not. Extended Data Figure 1-1 describes the most important parameters modulated for datasets in each of the three parameter regimes, “Unphysiological”, “Canonical”, and “Physiological”, along with the False Positives and False Negatives, for each of the ten implemented algorithms.

Time-series fluorescence data for various cells was extracted using Suite2P (Pachitariu et al. 2017). All further analysis and code development was done on MATLAB R2017b and batch analysis runs were performed on MATLAB R2021a. The average of the fluorescence values for cell specific pixels is then converted into the fold change relative to the baseline (dF/F0; F0 as 10th percentile), for every marked cell, in every trial (Figure 1B). These dF/F traces were used for the rest of the analysis.

### Curating a library of calcium events

For all synthetic data experiments, we used one good quality 2-P recording session’s worth of data from one animal. We mapped our imaging dataset into a matrix of dF/F values for all cells, trials, and frames. We then identified calcium events as signal deviations that were above a threshold (mean + 2*standard deviation) for more than 4 consecutive frames (frame rate: 14.5 Hz or ~70 ms per frame). Once identified, we curated a library for each event by a cell, and saved the respective start indices and widths. Using this library we generated synthetic data by inserting experimental calcium events into the time-series for each simulated cell. This approach just uses a time-series of signal bins and amplitudes, hence is signal-agnostic and could be applied to other imaging and recording modalities. In the interests of data integrity, our synthetic datasets were watermarked to be distinguishable from real physiology datasets.

### Generating Synthetic Data

Synthetic data was generated using a custom-written MATLAB function script “generateSyntheticData()” in the provided code repository. We preallocated and set up a 3-D matrix of zeros (as cells, trials, frames), and added calcium events sampled from the Calcium Event Library at frames (time) determined by the synthesis algorithm. The input parameters to this algorithm included timing, noise, imprecision, event width selection, hit-trial ratio, background activity, and several others. We aimed to cover the most likely conditions to affect timing and other experiment design properties. In more detail, we generated synthetic datasets using the following control parameters:

- Time Cell Percent Value: Number between 0 and 100. This sets the number of cells that are assigned tuned calcium activity as a percentage of total cells, and controls the number of Positive and Negative Class cells in the dataset.
- Cell Order Value: ‘basic’ or ‘random’. In ‘basic’ mode, Time Cells are indexed lower than Other Cells. In ‘random’ mode the indices of Time Cells and Other cells are randomly selected. This should have no impact on algorithm detection but is useful for visualisation.
- Max Hit Trial Percent Value: Number between 0 and 100. This sets the maximum possible fraction of total trials, during which a Time Cell will exhibit tuned calcium activity.
- Hit Trial Percent Assignment Value: ‘fixed’ or “random”. In ‘fixed’ mode, the number of Hit Trials is set as defined by Max Hit Trial Percent. In ‘random’ mode, the number of Hit Trials is calculated by randomly picking a value from a range (½*Max Hit Trials, Max Hit Trials).
- Trial Order Value: ‘basic’ or ‘random’. In ‘basic’ mode the Hit Trials are indexed lower than Miss Trials. In ‘random’ mode the indices of Hit and Miss Trials are randomly selected. Specific patterns of Hit and Miss Trials for a session have not been reported in physiology, so this feature is not implemented.
- Event Width Value: {0-100 percentile value, Integer N}. For each cell, this defines the selection of events based on width in frames. The percentile value is estimated from the histogram of all event widths. The variance of this selection is set by “N”, which adds N*Std. Dev. to the selection. All synthetic cells exhibit a range of different calcium events. This is considered an important parameter.
- Event Amplification Factor Value: Number from 0 to +∞. This allows additional control to multiplicatively amplify any chosen calcium event, before incorporation. Our library was curated from physiologically recorded signals. The default value is 1.
- Event Timing Value: ‘sequential’ or ‘random’. In ‘sequential’ mode the time of peak calcium activity is reflected by the indexing of the Time Cells. In ‘random’ mode the time of peak calcium activity is randomly dispersed over the trial frame-points.
- Start Frame Value: Number from 0 to total number of frames. This sets the timing of the first cell in a Time Cell Sequence.
- End Frame Value: Number from 0 to total number of frames. This sets the timing of the last cell in a Time Cell Sequence.
- Imprecision FWHM Value: Number from 0 to total number of frames. This sets the lower and upper bounds for the difference in timing of calcium activity across trial pairs for a Time Cell. We use this parameter to model trial to trial variability and is considered an important parameter to test.
- Imprecision Type Value: ‘none’, ‘uniform’ or ‘normal’. In ‘uniform’ and ‘normal’ modes, the trial pair Imprecision is picked from a Normal and Uniform Distribution, respectively. In ‘none’ mode, the trial pair Imprecision defaults to 0.
- Noise Value: ‘gaussian’ or ‘none’. In ‘gaussian’ mode the noise is sampled as a time-series vector with points from a Gaussian distribution. In ‘none’ mode, the Noise Percent defaults to 0.
- Noise Percent Value: Number from 0 to 100. This allows scaling for any sample noise point, based on the max signal value for any cell.
- Add Background Spikes for Time Cells Value: Boolean 0 or 1. This switch controls the incorporation of background (untuned) activity for putative time cells.
- Add Background Spikes for Other Cells Value: Boolean 0 or 1. This switch controls the incorporation of background (untuned) activity for other (non-time) cells.
- Background Distribution Mean Value: Number from 0 to +∞. This sets the Mean (λ) of the Poisson Distribution to sample from when selecting how many background events to add per trial, for any given cell.

### Implementation of a Reference Quality measure, Q

In order to compare the readouts from the various time-cell detection methods, we implemented a reference measure of Quality (Q) of synthetic time cells that utilized the known inputs to the generation algorithm.

Based on preliminary analysis, we selected following five parameters as the most likely to affect the behaviour and detection of Time Cells:

1. Noise
2. Event Width
3. Imprecision
4. Hit Trial Ratio
5. Background Activity

Accordingly, we were able to calculate a Reference Quality measure, using the following equation:

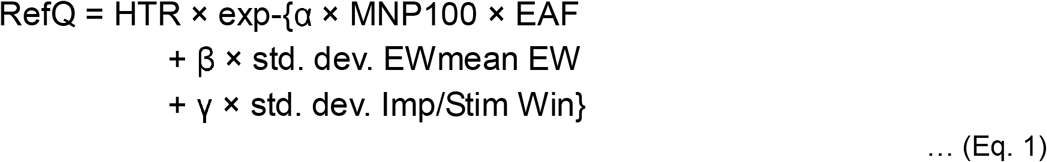

… where HTR: Hit Trial Ratio
MNP: Max Noise Percent (%)
EAF: Event Amplification Factor
EW: Event Widths (frames)
Imp: Imprecision (frames)
Stim Win: Stimulus Window (frames)
α: 1
β: 1
γ: 10

The values of α, β, and γ, were set to have comparable effects of each of the terms inside the exponent. This Reference Q was useful for debugging code and was the basis for a further metric for time-cell classification discussed below. A representative synthetic activity trace for ‘low’ and ‘high’ values of each of these five parameters is shown in Figure 1. All modulations for the datasets in this study along with the estimates for False Positives and False Negatives, across all algorithms are shown in Figure 1-1.

### Separate analysis modules were developed for three categories of analysis

We implemented three analysis modules: *ti*, *r2b* and *peq*, shorthand for temporal information, ridge-to-background, and parametric equations. The ti module implements three algorithms from Mau et al., 2018. The r2b module implements two algorithms from Modi et al., 2014. The peq module computes estimates for noise, hit-trial ratio, event width and imprecision, and estimates a Q score as above. All three methods were implemented in C++ with a PyBind11 interface to Python. This combination is fast and efficient in memory use, and also has the ease-of-use of Python. Thanks to the native MATLAB interface to Python, all three methods can also be called from MATLAB.

### Synthetic Datasets generated and analysed in batch mode

We generated datasets pertaining to Parameter Sensitivity Analysis by modulating one of the four main parameters and setting the others to non-interference levels. In this manner we devised 99 cases to study in which one of the main parameters was varied. Note that in these cases the resultant activity was in an un-physiological regime because other sources of variation were kept to low levels so as not to interfere with the parameter of interest. With 3 randomised shuffles, we generated 297 unique datasets.

We wanted to use more realistic datasets, where we would modulate one of the four parameters while keeping the others to ranges typical of physiological data. We devised 12 canonical cases. With 10 randomised shuffles each, we generated 120 additional unique datasets in the canonical regime. Finally, we devised 12 physiological regime cases, identical to those in the canonical regime, with the addition of background (untuned) activity. This yieled another 150 datasets, with randomization.

Altogether, we had 567 unique datasets for our tests, each with 135 cells (total: 76545 cells), 60 trials, and 246 frames/trial. Except when the percent time cells were modulated, all datasets featured 50% time cells.

We next implemented an analysis pipeline to run all the datasets through the time-cell detection algorithms, yielding scores and predictions for each case. Finally all the scores and predictions were collated for comparison and benchmarks as shown in the schematic (Figure 2).

**Figure 2:**
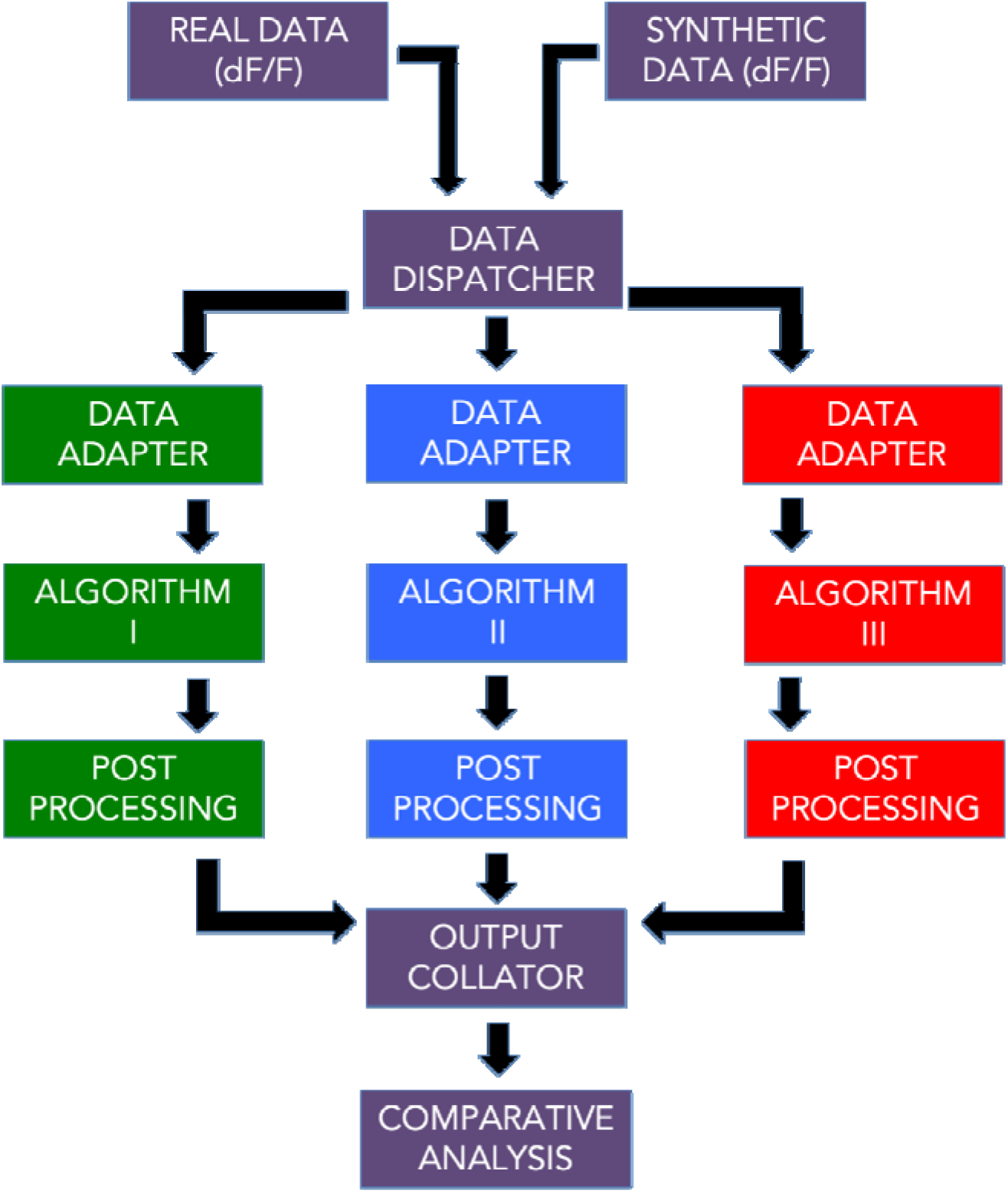
A schematic representation of the analysis pipeline. Physiology data as well as synthetic data was analysed by 10 different implemented algorithms and the output was collated for comparative benchmarks.

### Metrics for time cell Classification Performance

Recall is inversely proportional to the number of False Negatives (Type II error) and is the fraction of True Positive Class predictions over all Positive Class ground labels.

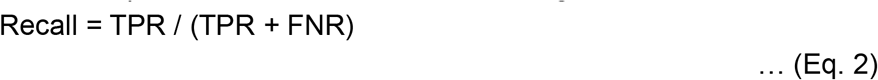

Precision is inversely proportional to the number of False Positives (Type I error) and is the fraction of True Positive Class predictions over all Positive Class predictions.

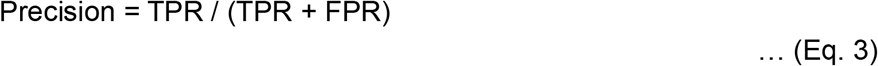

F1 Score is the harmonic mean of Recall and Precision.

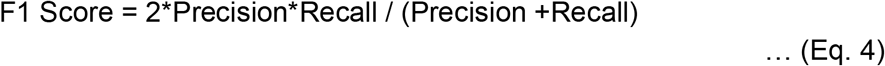

… where,
TPR: True Positive Rate
FNR: False Negative Rate
FPR: False Positive Rate

Here are the definitions for predictive/classification performance evaluation (Table 1).

**Table 1:**
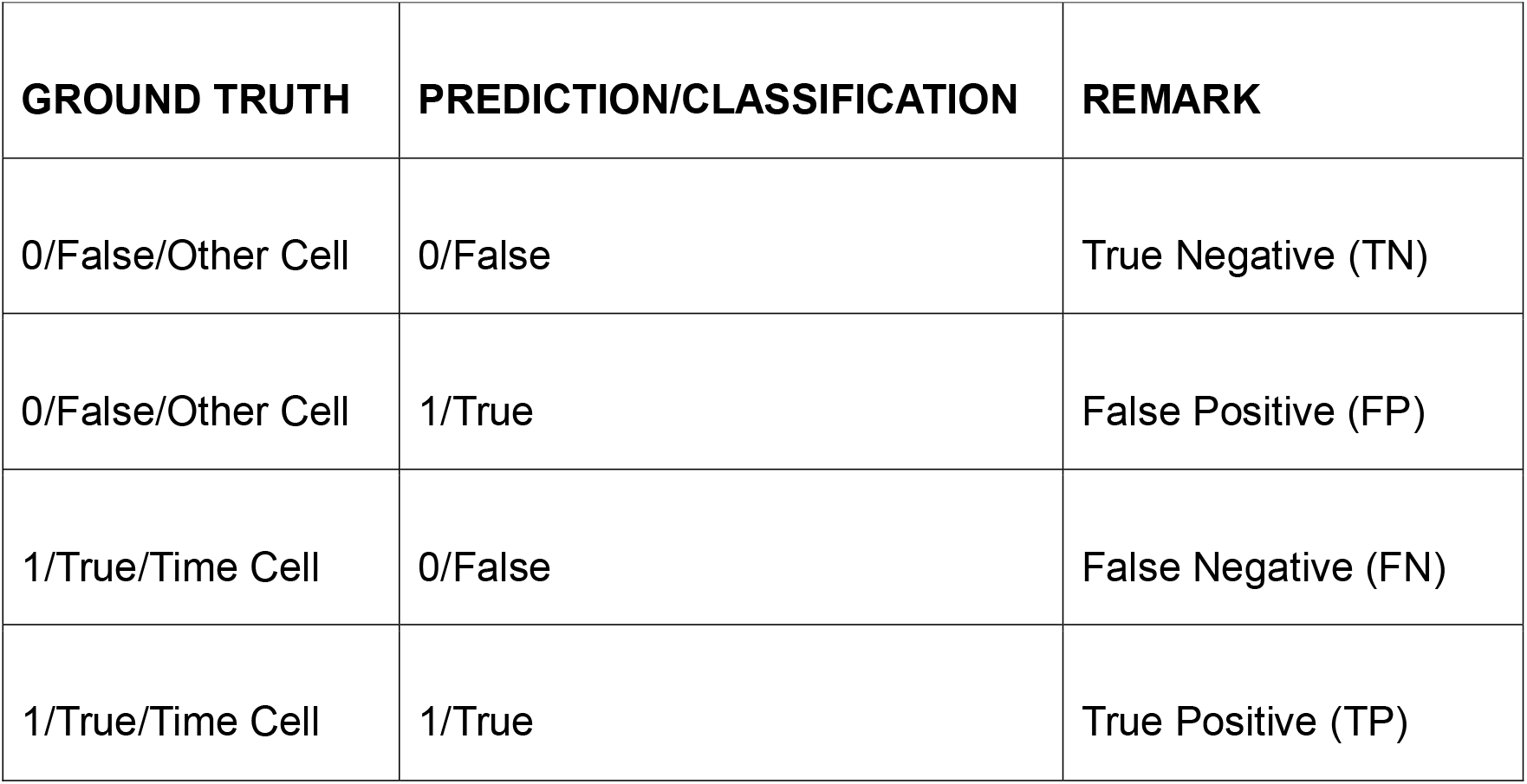
Definitions for predictive/classification performance evaluation. For each detection algorithm, the classification results were compared to known ground truth values to get the total number of True Positive (TP), True Negative (TN), False Positive (FP), and False Negative (FN) cases.

| GROUND TRUTH | PREDICTION/CLASSIFICATION | REMARK |
| --- | --- | --- |
| 0/False/Other Cell | 0/False | True Negative (TN) |
| 0/False/Other Cell | 1/True | False Positive (FP) |
| 1/True/Time Cell | 0/False | False Negative (FN) |
| 1/True/Time Cell | 1/True | True Positive (TP) |

## Results

We developed a pipeline (Figure 2) with ten different algorithm implementations for Time Cell detection, which involve scoring and then classifying cells.

Here, we describe the implementation of each of the methods.

### Time Cell Scoring Methods and Classification

#### Temporal Information – *tiBoot, tiMean, tiBoth, tiMean-O, tiBase-O* (Mau et al., 2018)

Here we used the algorithm from Mau et al, as follows. There was an initial criterion of cells to have activity in at least 25% of trials. Their activity was summed into event time histograms with a bin size of 3 frames. The Temporal Information (TI) was estimated using Eq 5,

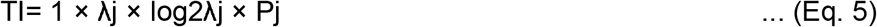

… where, λ is the Average Transient rate for each cell λj is the Average transient rate for each cell in bin “j” Pj is the Probability that the mouse was in time bin “j”

Bootstrapping was used to determine if each cell had a TI greater than chance. We circularly randomized the frame points to develop a random activity model (1000 iterations) and classified cells as time cells if λ > λrand in greater than 99% of the models for at least 2 consecutive bins. We implemented the activity filter from Mau et al. 2018, by considering the trial-averaged peak of the calcium traces for each of the cells, and testing for significance using bootstrapping (*tiMean*). A logical AND operation between the prediction lists for tiBoot and tiMean, provided us with the full Mau et al., 2018 Temporal Information based detection algorithm (*tiBoth*).

Additionally, we used Otsu’s Threshold (Otsu, 1979) on the Temporal Information scores as well as the trial-averaged peaks to get *tiBase-O* and *tiMean-O* using the MATLAB function “graythresh()” (https://in.mathworks.com/help/images/ref/graythresh.html). The purpose of adding the Otsu’s threshold based classification step was to study how well the scores could be classified with a fast thresholding method, rather than the computationally expensive bootstrap.

#### Ratio of Ridge/Background – *r2bMean, r2bBoot, r2bBase-O* (Modi et al., 2014)

Here we re-implemented the algorithm from Modi et al. 2014. The time of peak response for each cell was identified in averaged, non-overlapping trials’ worth of ΔF/F traces, in the CS-onset to US-onset period, or as specified. The rest of the trials were averaged and the summed area under the time of peak was estimated. The ridge was then defined to be a 200 ms window centered at the peak. Next, we calculated the summed area in the ridge window as well as the background (non-ridge frames) to get the ridge to background ratio. As a control condition, these traces were given random time-offsets and then averaged. An independent time of peak was identified for each random-offset, averaged trace and ridge to background ratio calculated for it. This bootstrapping was repeated 1000 times for each cell’s data and averaged. The reliability score was then calculated individually, for each cell, as the ratio of the ridge to background ratio for aligned traces to the mean of that of the random-offset traces (*r2bMean*).

We also studied the significance of each cell’s raw r2b values by comparing them to each of the r2b values of the randomized datasets, thresholding significance at the 99th percentile (*r2bBoot*). Finally, the raw r2b values were also thresholding using Otsu’s Thresholding (*r2bBase-O;* Otsu 197).

#### Parametric Equations - *peqBase* and *peqBase-O* (in-house)

We developed this method to score cells in a manner similar to the Reference Quality, which uses the known ground truth of the input parameters given to the generator functions for the synthetic dataset. Rather than using the known inputs, this method computes the corresponding parameters read out or estimated from the dataset, whether synthetic or real. It is applicable to labelled or unlabeled datasets. It is defined as:

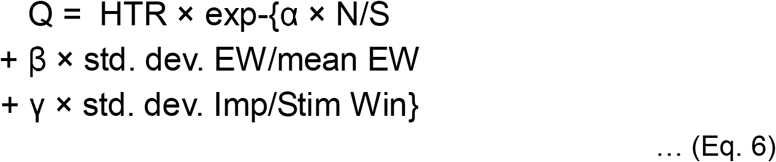

… where HTR: Hit Trial Ratio
N/S: Estimated Noise/Signal
EW: Read out Event Widths (frames)
Imp: Estimated Imprecision (frames)
Stim Wind: Stimulus Window (frames)
α: 10
β: 1
γ: 10

While 10xα was required, β, and γ, were inspired by the same used for Reference Q. Classification was then performed using Bootstrapping (as described above) as well as Otsu’s Threshold.

All of these implemented algorithms can handle unlabelled (real) or ground truth labelled (synthetic) data.

A schematic to describe the steps involved in each algorithm is shown (Figure 3). We were then able to run all our synthetically generated datasets through each of the ten implemented algorithms and perform comparative benchmarks.

**Figure 3:**
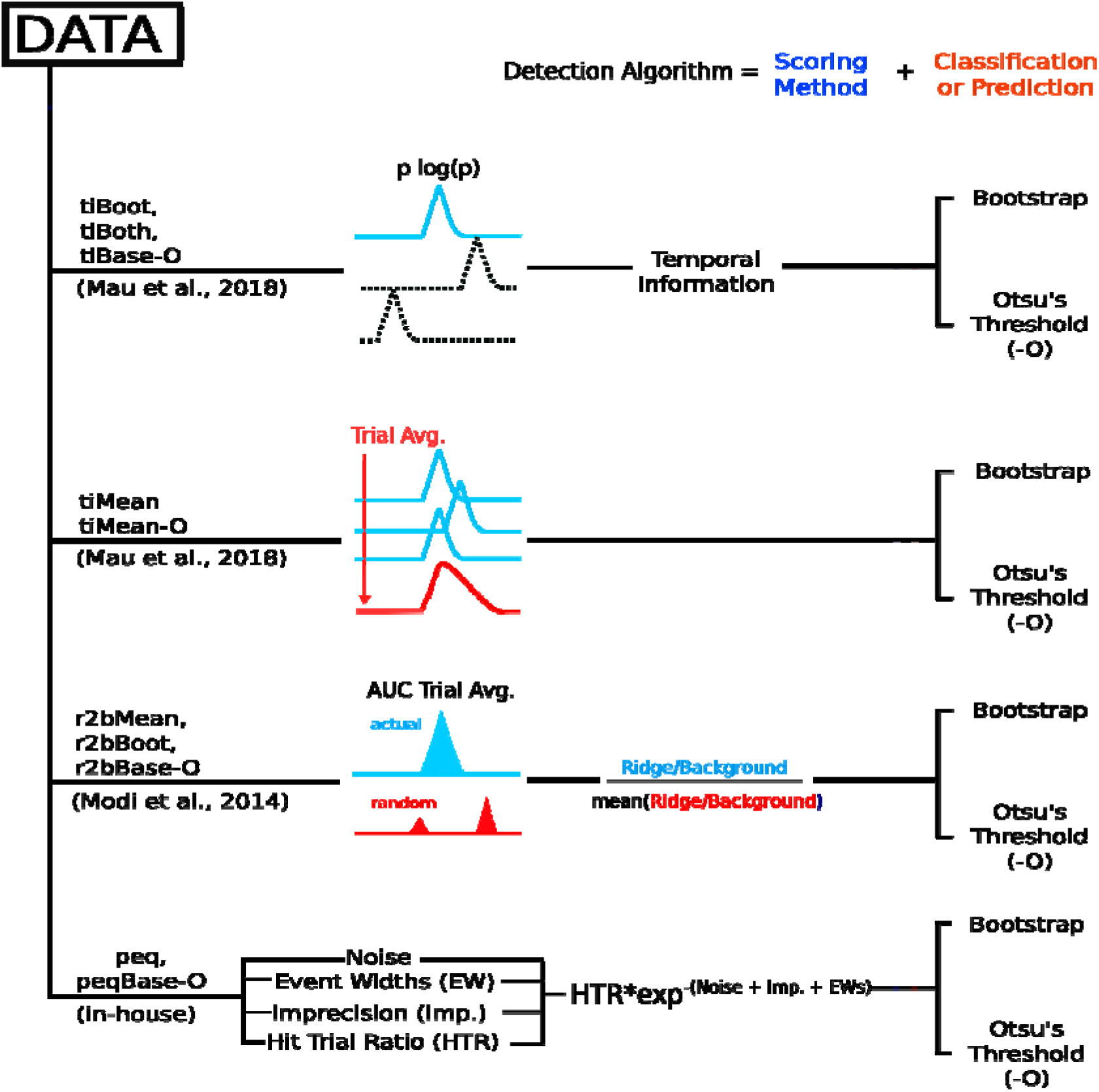
Schematic representation of the implemented algorithms, involving four different scoring methods followed by a classification step (Bootstrapping or Otsu’s automatic threshold) to have ten complete time cell detection algorithms.

### Good predictive power in time cell quality scores despite different distributions

We ran each of the analysis methods on our synthetic datasets to assess how they scored the (known) time cells. There were four methods that provided a scoring function for time-cell classification: *tiMean, tiBase, r2bBase* and *peqBase* (Figure 4A–4D). By inspection, these methods appeared to have distinct distributions. Below we describe how we compare the distributions using correlation analysis. In subsequent sections we describe other methods in our study that used these scores to generate a categorization through thresholding or bootstrap.

**Figure 4:**
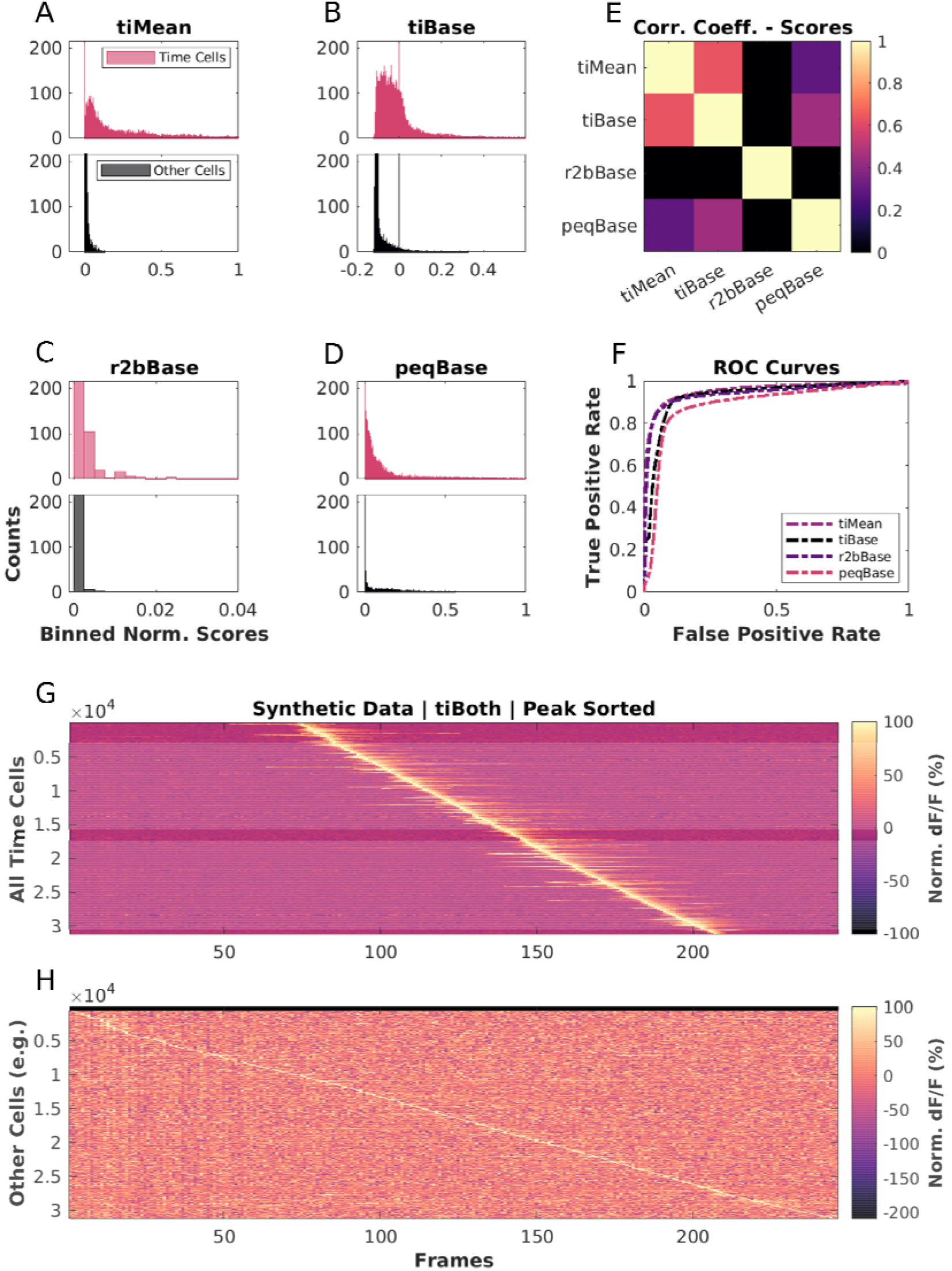
Base scores for different methods differ in their distributions but all have good predictive power. Scores for top (blue): time cells; bottom (red): other cells, across A: *tiMean*, B: *tiBase*, C: *r2bBase*, D: *peqBase* D, E: Pairwise Correlation Coefficients between the distributions of analog scores (pooling time cells and other cells) by each of the four scoring methods, F: Receiver-Operator Characteristic (ROC) Curves after Generalized Linear Regression using the respective distributions of scores and comparisons with known Ground Truth G-H: Trial-Averaged calcium activity traces for cells classified as G: time cells, H: other cells.

In these synthetic experiments, time cells were generated with a single calcium event per Hit Trial. Event insertions into the synthetic datasets were subject to Noise, variable selection of event widths, trial-pair or timing Imprecision, and Hit Trial Ratio. We generated 99 unique un-physiological combinations (3x randomised shuffles) 12 unique canonical regime combinations (10x randomized shuffles), as well as 15 unique physiological regime combinations featuring background activity (10x randomised shuffles). In all, we performed our comparative analysis studies using 567 datasets, each with 135 cells, 60 trials/session, and 246 frames/trial at 14.5 Hz). We found that only TImean and TIbase had a correlation coefficient of ~0.6, whereas other pairs were correlated below 0.4 (Figure 4E). Generalised Linear Regression (GLM) models were generated to look for the ideal thresholding value for the best classification predictions by each method. We used the MATLAB implementation of GLMs (fitglm(), https://in.mathworks.com/help/stats/fitglm.html). This is a linear model assuming a binomial distribution of categories (0 or 1, i.e., other cell or time cell; (Collett, 2002). We obtained good predictive power for the four methods that provided a scoring function for time-cell classification. We generated ROC curves by going over the full range of thresholds for the range of scores for each method (ROC Curves, Figure 4F). We found that each distribution of scores had good predictive power, since ideal thresholds could be found to maximize TPR/FPR in all cases. We used the TI-Both categorization to distinguish time cells (Figure 4G) from other cells (Figure 4H), and plotted Trial-Averaged calcium traces to visually assess quality of classification as seen from raw data. Overall, each of our methods had distinct distributions of their base scores, but all had good predictive power for classification. The outcome of the classification steps is described in the next sections.

### All algorithms exhibit near perfect Precision with good Recall

Next, we used the scores to classify the cells in our synthetic datasets, compared the predictions to ground truth, and established summaries for True and False cases. Confusion matrices were estimated to compare the predictions (classifications) for each algorithm, with reference to ground truth, and are shown (Figure 5A and 5B). All methods exhibit very good Precision (true positive classifications over the sum of all positive classifications), suggesting low False Positive Rates (Type I error, Figure 5C). Most algorithms also generate good values for Recall (true positive classifications over ground-truth positives). We observed F1 Scores (Harmonic mean of Recall and Precision) greater than 0.75, all the way to 1 (perfect score), for most of the algorithms, as shown (Figure 5C), suggesting overall usability.

**Figure 5:**
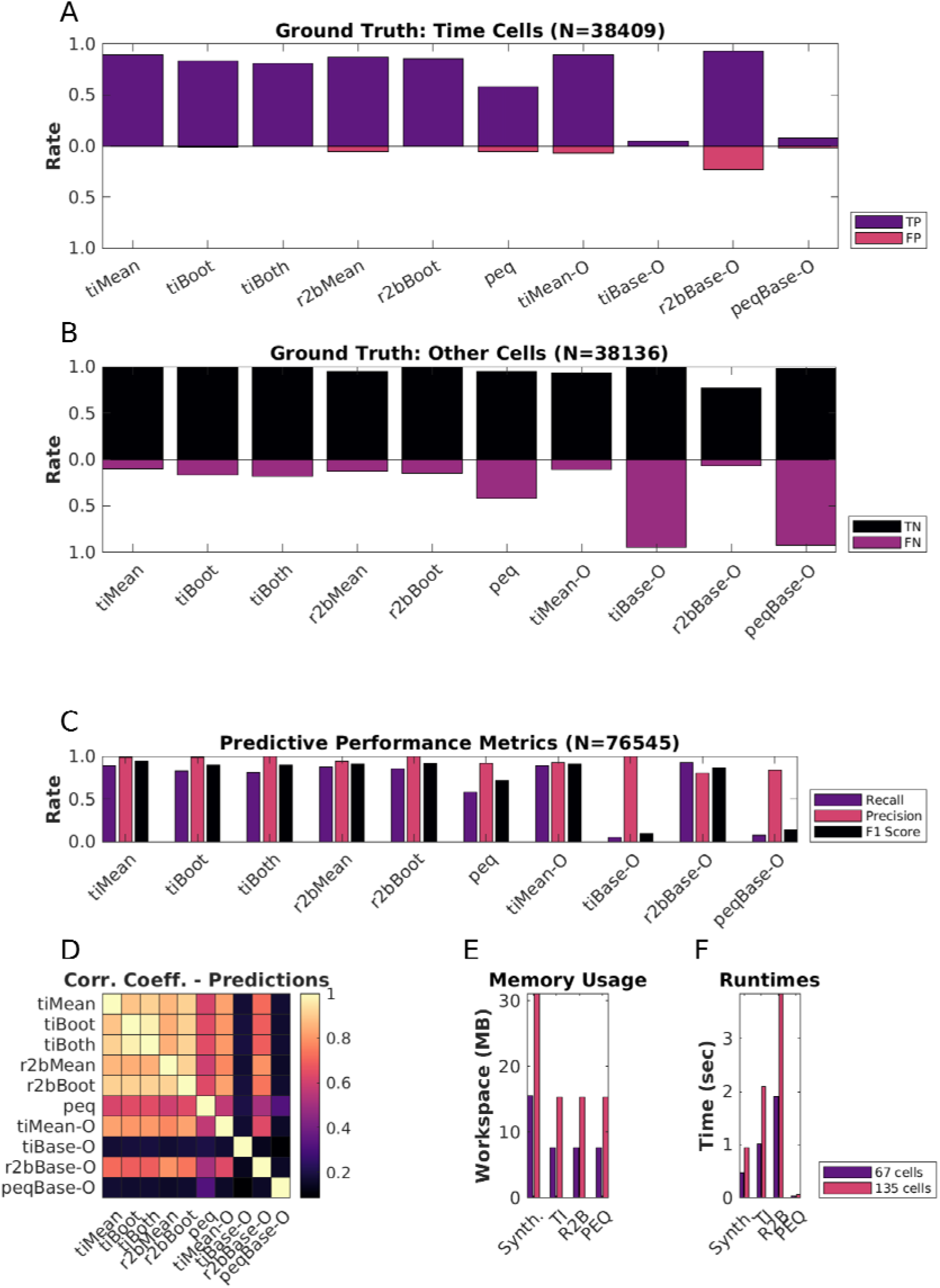
Good predictive performance by all algorithms. A-B: Classification performance of each of the ten implemented detection algorithms. A: True Positives (TP; maroon), False Positives (FP; red).B: True Negatives (TN; black), False Negatives (FN; maroon). C: Predictive Performance metrics (Recall=TP/(TP+FN), Precision=TP/(TP+FP), and F1 Score=Harmonic mean of Recall and Precision) to consolidate the confusion matrices. D: Pairwise Correlation Coefficients between the boolean prediction lists by each of the ten detection algorithms. Note that the first six methods correlate strongly. E: Average Memory usage per dataset by the implemented algorithms on datasets with either 67 cells (blue) or 135 cells (red). F: Average runtimes per dataset by the implemented algorithms on datasets with either 67 cells (blue), or 135 cells (red).

We noticed moderate to strong correlation (>0.8) between the boolean prediction lists for *tiMean, tiBoot, tiBoth, r2bMean* and *r2bBoot* (Figure 5D), but only weak to moderate correlation (<0.6) between the other pairs of predictions. The tiMean-Otsu method does slightly better (correlation ~0.7 with the first 5 methods).

### Algorithms differ in memory use and speed

Hardware and runtime requirements are a secondary, but practical concern when designing analysis of large datasets, and are specially relevant for experiment designs that require online analysis. We therefore looked at how memory use and runtime scaled on a per dataset basis when considering 67 or 135 cells per dataset (2x).

We ran the memory usage and runtime experiments on a gaming laptop (Lenovo Ideapad 3 Gaming) with a 6 core AMD Ryzen 5 4600H, 16 GB DDR4 RAM (3200 MHz) running MATLAB R2021a on Ubuntu 20.04. Note, however, that we have implemented all the time-cell algorithms in serial and these do not use the additional cores. We found that most algorithms ran to completion requiring ~15 MB/dataset at a rate of ~1-4 sec/dataset (135 cells/dataset). With 67 cells/dataset, the memory requirement and runtimes are approximately halved, suggesting that computational costs in memory and time were roughly linear with dataset size. We note that the analysis algorithms work independently for each cell. Thus in principle the analysis could be run in an embarrassingly parallel manner and should scale well on multi-core architectures.

The synthesis of the main benchmarking datasets (N = 567 datasets or 76545 total cells) required a more powerful analysis machine, running a 6 core AMD Ryzen 5 3600, 32GB of DDR4 RAM, running MATLAB R2021a on Ubuntu 20.04. Dataset batches up to ~30 datasets (N = 40500 cells), however, could be easily handled by a less powerful laptop. The memory usage and runtime for 135 cells per dataset were accordingly, ~30 MB/dataset requiring ~1 sec to complete. Thus, the methods scale readily to handle large datasets on modern hardware.

### Physiological range tests show sensitivity to Noise but not to other features of the dataset

We next set out to see how these methods would work in estimated physiological ranges of signal confounds. Given our categorical labels on the synthetic data, we were able to split the datasets to look for the effects of the five main parameters, Noise, Event Widths, Imprecision, Hit Trial Ratio, and Background Activity. We first computed the baseline physiology readouts keeping Noise to 10%, Event Widths to the 60th percentile (+/− 1 std. dev.), Imprecision to 0 frames, Hit Trial Ratios to a range of 33-66%, and background activity to 0.9-1.2 events/trial for Time Cells (~50% of all synthetically generated cells, N = 50 baseline datasets, 135 cells/dataset, 60 trials/dataset). Next, we established dependency slopes for each of the algorithms, based on their predictions (N = 10 randomised shuffles for each case, Figure 6B to 6F, Figure 6–1, Figure 6–2).

**Figure 6:**
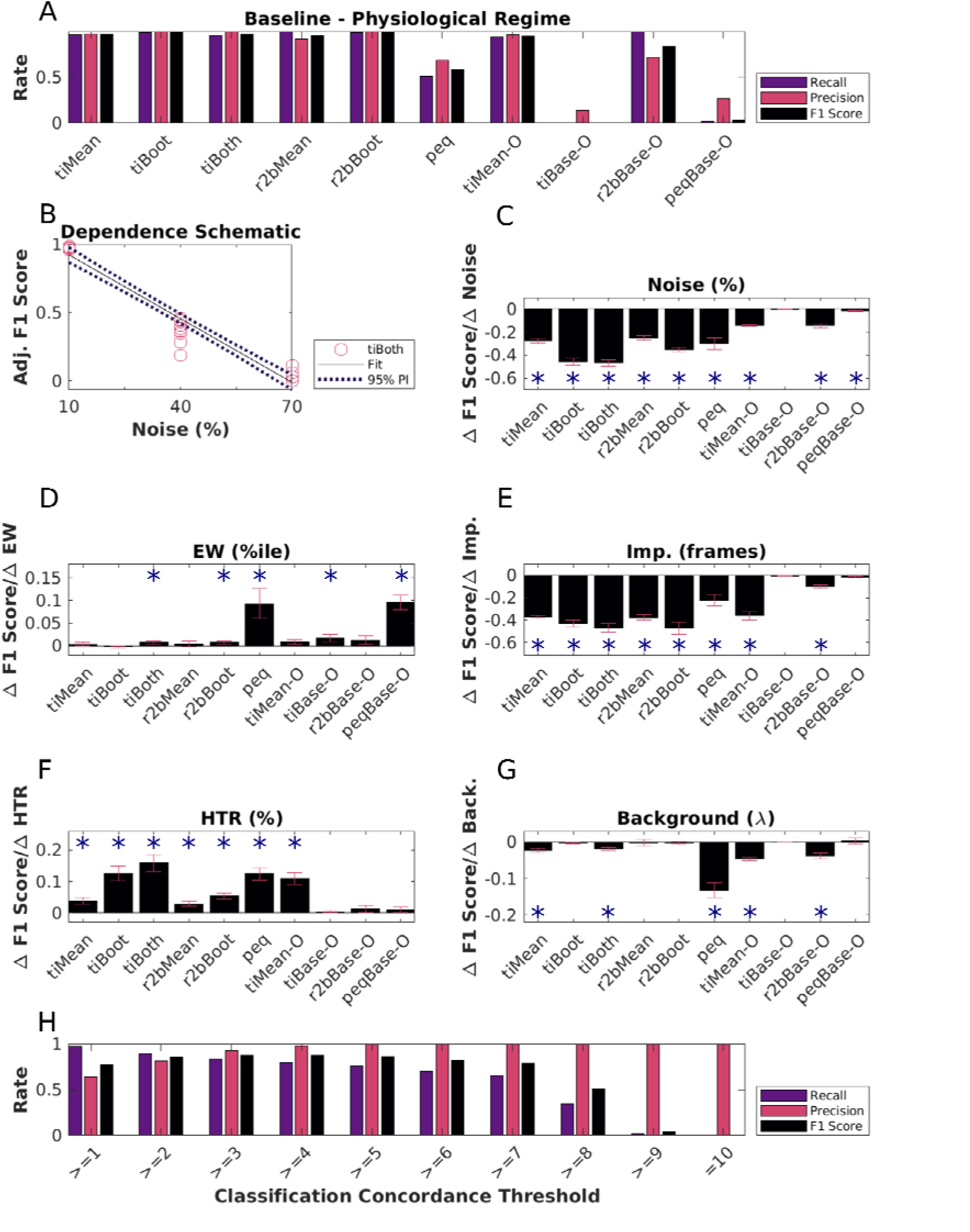
Physiological Sensitivity Analysis and Concordance. A: Classification performance scores for all algorithms with the baseline Physiology synthetic datasets (N = 6750 cells). The first five methods perform well. Peq does poorly by all measures when confronted with physiology-range activity variability. Otsu’s threshold method for score classification also does not work well for any method under physiological conditions. B. Dependence of F1 score on noise as a schematic. This has an overall negative slope (dashed line) which was used for panel C, TI-both. A similar calculation was performed for each method. Panels C-G: Parameters were systematically modulated one at a time with respect to baseline and the impact on classification score for each algorithm was estimated by computing the slope, using repeats over 10 datasets each with an independent random seed. Significant dependence on the perturbing parameter was determined by testing if the slope differed from 0 at p<0.01, indicated by asterisks using the MATLAB function coefTest(). Plotted here are bar graphs with mean and error as RMSE normalized by the square root of N (N = 10 datasets). C: Dependence on Noise %, D: Dependence on Event Width percentiles, E: Dependence on Imprecision frames, F: Dependence on Hit Trial Ratio (HTR; %), and G: Background Activity (Poisson Distribution mean, λ). H: Classification performance using Concordance for a range of classification thresholds. Extended Data Figure 6-1 describes the three-point line plot dependency curves for the F1 Score for each of the implemented algorithms against each of the five main parameters modulated, as the mean of N=10 datasets for each case, with error bars as standard deviation. Extended Data Figure 6-2 showcases the linear regression fits for the same, with 95% prediction intervals (PI), used to estimate the slopes of the various dependency curves.

Most methods exhibited a negative dependence of Noise (range: 10% to 70%) on prediction F1 Score (Figure 6B). Although many methods are designed with some form of denoising strategy (trial-averaging, etc.), as expected all algorithms ran into classification difficulties at higher Noise levels. This reinforces the value of relatively high signal to noise recordings.

The relative insensitivity to event widths (Figure 6C) is potentially useful for Calcium Imaging datasets where events may be slow, and in cases where slower tuning curves are expected. However, this criterion may need to be stringent for analyses that need to precisely identify fine differences in cell responses.

We observed that most algorithms were insensitive to how frequently time cells were active across trials in a session (HTR). This is possibly the reason for the potential confusion amongst physiologists with regard to how many time cells were expected in a recorded dataset.

We found that the first six algorithms (*tiMean, tiBoot, tiBoth, r2bMean, r2bBoot*, and *peq*) gave equivalent predictions in ~66% of cases (Figure 6–1A). Next, we considered the various prediction lists across these top six algorithms and looked for consensus in Time Cell predictions from the most lenient threshold (“>=1” algorithm), incrementally through the most stringent threshold (“= 10” algorithms). We thus established a Concordance based metric for time-cell classification. We tested the predictive power of this Concordance based metric, which considers time cells based on consensus amongst the predictions from all the ten implemented algorithms. We identified differences in the classification performance, across the full range of concordance thresholds (Figure 6H). With lower threshold values (“>=4” and below), we notice a slight drop in the Precision, indicating an increase in False Positive Rate (Type I error). On the other hand, with increasing threshold values it is the Recall that drops, suggesting a higher False Negative Rate (Type II error). We find that a Concordance threshold of “>=4” achieves the best Recall, Precision, and F1 Scores, for Time Cell prediction (Figure 6F). The utility of this approach is subject to the availability of resources to apply multiple algorithms to each dataset.

### Time cells identified in Real Physiology recordings

We used the ten different implemented algorithms on in-vivo 2-P calcium recordings (N = 13 datasets, viz., 1759 isolated cells from 3 animals across chronically recorded datasets), to compare time-cell classification between the algorithms. As we observed for the synthetic data, experimental 2-P Ca traces also yielded different base scores from the four different methods (Figure 7A to 7D) Again, consistent with the synthetic data, the pairwise correlation was weak to moderate (Figure 7E). When we consider the boolean prediction lists (Figure 7F), we observed moderate pairwise correlation between tiMean, tiBoot, *tiBoth, r2bMean*, and *r2bBoot* (>0.5), and low or weak correlation between the other pairs (<0.5). This was consistent with observations for the synthetic data but the correlations were overall slightly weaker. The total number of Time Cells predicted were also different across the implemented algorithms (Figure 7G). Algorithms such as *r2bBase-O* and *peq*, which had more false positives (Figure 5B) also had more cells classified as time cells. The converse was not true. *r2bMean*, which had moderate false negatives as well as false positives on the synthetic dataset, classified very few of the experimental set as time cells. The Trial-Averaged activity of the detected time cells (Figure 7H; including False Positives) and other cells (Figure 7I), based on the predictions by tiBoth, are shown. The experimentally recorded time cells exhibited a characteristic widening of tuning curves (Pastalkova et al., 2008; MacDonald et al., 2011; MacDonald et al., 2013; Kraus et al., 2013; Mau et al. 2018) with tuning to later time points (Figure 7H).

**Figure 7:**
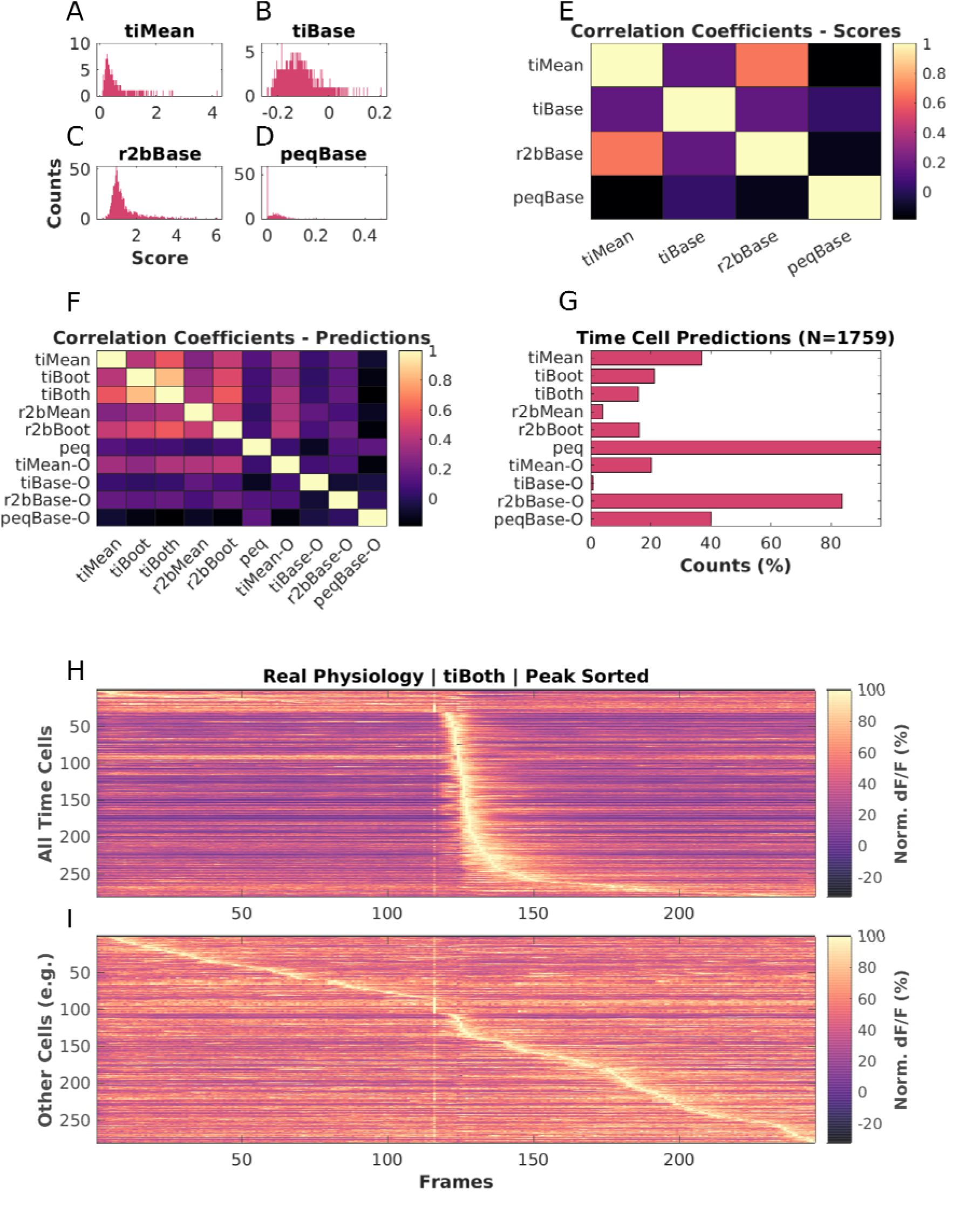
Analysis of experimental 2-P recordings of Ca2+ signals. A-D: Histograms of scores for physiologically recorded *in vivo* calcium activity from Hippocampal CA1 cells (Total N = 1759), by A: tiMean, B: tiBase, C: r2bBase, and D: peqBase. E: Pairwise Correlation Coefficients between the distributions of analog scores by the four scoring methods. F: Pairwise Correlation Coefficients between the boolean prediction lists by the ten detection algorithms. G: Numbers of Positive Class (time cell) predictions by each of the detection algorithms. H-I: Trial-Averaged calcium activity traces for H: time cells, and I: other cells. LED conditioned stimulus (CS) is presented at frame number 116, as seen by the bright band of the stimulus artifact. Most cells classified as time cells are active just after the stimulus. There is a characteristic broadening of the activity peak for classified time cells at longer intervals after the stimulus. Some of the cells at the top of panel H may be false positives because their tuning curve is very wide or because of picking up the stimulus transient. Similarly some of the cells in the middle of panel I may be false negatives due to stringent cutoffs, even though they appear to be responsive to the stimulus.

Overall, four of the algorithms from the literature seemed consistent in their classifications as well as having reasonable numbers of classified time cells. These were the three algorithms from Mau et al., 2018 (*tiMean, tiBoot*, and *tiBoth*), and the *r2bBoot* method derived from Modi et al., 2014. This is broadly in agreement with their performance on the synthetic datasets.

## Discussion

We have developed a full pipeline for comparing time-cell detection algorithms. This starts with synthetic datasets for benchmarking, in which we program in the ground truth of cell identity and timed activity, and a range of perturbations characteristic of experiments. These include Noise, Event Widths, trial-pair timing Imprecision, Hit Trial Ratio, and Background Activity. This resource is, in itself, a key outcome of the current study, and though it is designed for 2-P Calcium imaging data it can be extended to rate-averaged single-unit recordings. We built a pipeline for running and comparing the outcome from five methods derived from two previous studies, and one from the current work. These algorithms were applied to synthetic and experimental datasets and compared against each other and, where possible, against ground truth. We observed that most algorithms perform well and substantially agree in their time-cell classification, but there were different degrees of sensitivity to different forms of signal variability, notably Noise and Imprecision.

### The value of synthetic data in experimental science

Synthetic neural activity datasets are valuable in at least two main ways: evaluating algorithms for detection of important activity features, and for delivering stimuli to in-vitro and simulated neurons, so as to provide a more physiological context in which to study input-output properties (Abbasi et al., 2020). While we have deployed our synthetic dataset for the specific purpose of comparing time-cell detection algorithms, we suggest that it could also be useful for evaluating sequence analysis algorithms (Foster & Wilson, 2006; Ikegaya et al., 2004; Villette et al., 2015). Beyond the domain of neuronal data analysis, such synthetic datasets act as a test-bed for critique and development of analysis algorithms meant for deployment on real-world or typical use case data. They have been used previously to benchmark unsupervised outlier detection (Steinbuss & Bohm, 2020), explainable Machine Learning (Liu et al., 2021), Intrusion Detection Systems (Iannucci et al., 2017), 3D Reconstruction Algorithms (Koch et al., 2021), among several others. We report the first use of synthetic data pertaining to cellular physiology in the context of identifying time cells from network recordings. Moreover, our experiments study important operational differences across several previously published and new detection algorithms. Our dataset may also be valuable for the second use case, stimulus delivery. There is a growing body of work on network detection of sequences (Csicsvari et al., 2007; Foster & Wilson, 2006; Ikegaya et al., 2004; Jadhav et al., 2012; Malvache et al., 2016; Villette et al., 2015) or even single-neuron sequence selectivity (Bhalla, 2017; Branco et al., 2010). More realistic input activity patterns with a range of physiological perturbations may be useful probes for such experimental and theoretical studies. Further, experimenter-defined neural activity inputs through optogenetic stimulation has already begun to utilize more complex temporal patterns than static or periodic illumination (Bhatia et al., 2021; Dhawale et al., 2010; Schrader et al., 2008). Our approaches to synthetic sequential neuronal activity generation may be useful to add more physiological dimensions to the sequential activity employed in such studies.

### Further dimensions of time-cell modulation

Our experiments allowed us to probe for parametric dependence systematically across published and new algorithms. We observed little or no dependence of the predictive performance (F1 Score) of the various algorithms to Event Widths, Hit Trial Ratios, and Background Activity. We did observe the F1 Scores for most algorithms to be negatively dependent on Noise and Imprecision. On the one hand, this is a useful outcome in that different methods yield similar time-cell classification. It is a limitation, however, if the network uses such response features for coding, since it means that these methods are insensitive to relevant response changes. Further potential coding dimensions were not explored. Thus several potential behavioural correlates of tuned cells (Ranck, 1973), could not be studied in our experiments. Such correlates include but are not limited to measurements of spatial navigation (O’Keefe & Dostrovsky, 1971; O’Keefe & Nadel, 1978; Wilson & McNaughton, 1993) and decision making (Csicsvari et al., 2007; Davidson et al., 2009; Diba & Buzsáki, 2007; Foster & Wilson, 2006; Gupta et al., 2010; Karlsson & Frank, 2009; MacDonald et al., 2013; Villette et al., 2015), as well as navigation across tone frequencies (Aronov & Tank, 2014). While each of these further inputs would be interesting to incorporate into synthetic datasets, this requires that the time-cell generation algorithm itself incorporate some form of simulation of the neural context. This is beyond the scope of the current study.

A specific limitation of our dataset is that it assumes that time is encoded by individual neurons. This leaves out population encoding schemes in which no one cell responds with the level of precision or consistency that would clear the criteria we use. For example, many of the same studies that utilise the methods tested here also use neural network decoders to report time (Mau et al. 2018). Such decoders might detect time encoding without time cells. A similar situation of individual vs. network coding appears for the closely related problem of sequence representation. Place cell replay sequences have been shown to be modulated by the prevalence of location specific aversive (Wu et al., 2017) as well as appetitive stimuli (Bhattarai et al., 2020). Such physiological findings have been the subject of theoretical models of behaviour planning (Foster, 2017; Mattar & Daw, 2018), and have been reported to improve performance on multiple Atari games by artificial neural networks (Mnih et al., 2015) featuring salience detection and experience mapping. We suggest that synthetic data for such higher-order encoding schemes might be a useful tool, and could draw upon the approaches in the current study.

### Comparative Analysis Benchmarks and Concordance

A particularly challenging time-cell classification problem is when the same cells may play different timing roles, such as forward and reverse replay. This is made more difficult because of the relative rarity of forward replay sequences over the more typical reverse replay (Diba & Buzsáki, 2007; Foster, 2017). Pre-play is also a topic of some debate (Dragoi & Tonegawa, 2013; Foster, 2017). At least one possible problem in such debates is the degree of consistency between time-cell or sequence classifiers. Our pipeline allows for a) error correction in case of non-concordant classifications, b) suggest candidate algorithms with a dependence on dataset features like Event Widths, Imprecision, and Hit Trial Ratio, as well as c) the possibility to expand the detection regime in more realistic physiological datasets using Concordance.

### Which algorithms to use?

We did not set out to rank algorithms, but our analysis does yield suggestions for possible use domains based on sensitivity to experimental perturbations (Figure 8). In cases where runtime and compute resource use is a concern, we recommend using the Temporal Information method with Bootstrap along with the activity filter (tiBoth). Combinations of *tiBoth* with *r2bBoot* may be useful where there are rare and potentially multimodally tuned time cells (Pastalkova et al. 2008; Villette et al. 2015), either to combine their classification for stringent time-cell identification, or to pool their classified cells. While it is tempting to use Otsu’s threshold as a very fast alternative to bootstrapping, we found that none of the Otsu variants of these methods did a good job of classification. Ultimately, five of our algorithms *tiMean, tiBoot, tiBoth, r2bMean* and *r2bBoot:* all based on either Mau et al., 2018 or Modi et al., 2014, have very good Precision, and classify with very few false positives (low Type I error). Many methods are susceptible to classification errors if the dataset has high noise.

**Figure 8:**
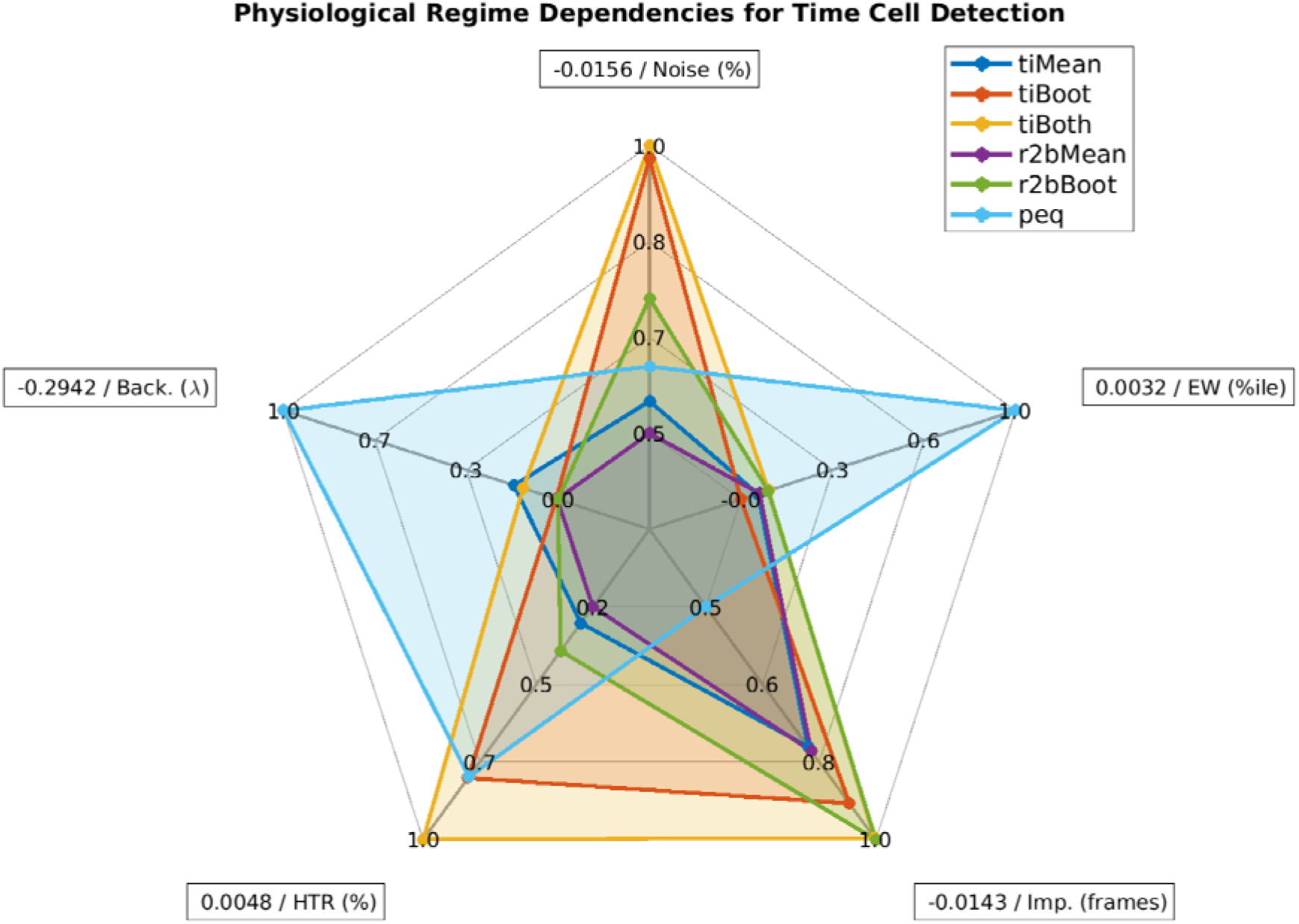
Spider Plot Summary: Relative sensitivity of the six best detection algorithms (tiMean, tiBoot, tiBoth, r2bMean, r2bBoot, and peq) to the five main parameters for data variability, Noise (%), Event Widths (%ile), Imprecision (frames), Hit Trial Ratio (%), and Background Activity (λ). A perfect algorithm would have very small values (i.e., low sensitivity) for each of the parameters, and thus occupy only the smallest pentagon in the middle. Note that even the maximal absolute value of sensitivity for most parameters (outer perimeter) is quite small, indicated in boxes at the points of the spider plot.

Here we also implemented the parametric equation (*peq*) algorithm. It is not very good for time-cell classification per se, as it isprone to false positives and is susceptible to noise and low hit-trial ratios. However, it generates useful additional estimates of the four key parameters of real data, namely: noise, hit-trial ratio, event width and imprecision. This is useful for a first-pass characterization of the properties of the dataset.

### Sequence detection in large-scale recordings and scaling of analysis runs

The discovery of replay over the past two decades, has benefitted from the technological advances made in increasing the cellular yield of network recordings and has been reviewed previously (Foster 2017). Further advances such as with the large scale recordings of ~10^3^ single units by electrical recording using Neuropixels (Jun et al., 2017), fast volumetric fluorescence scanning with upto ~10^4^ cells using Resonant Electro-Optic Imaging (Bowman & Kasevich, 2021; Pachitariu et al., 2016; Poort et al., 2015), ~10^3^ Mesoscopes (Sofroniew et al., 2016), as well as advances in automated cell ROI detection, denoising, and neuropill subtraction (Pachitariu et al., 2016; Pnevmatikakis et al., 2016) only increase the scale and size of datasets, likely leading to longer analysis runtimes. In addition to our recommendations above for the Temporal Information/Boot method for scalable time-cell analysis, our C++/Python implementations may also be useful in further optimising these methods. Our implementations allow for relatively fast analysis of the same datasets with multiple algorithms.

## Supporting information

All Supplementary Figures with Legends

## Code and Resource Availability

The code/software described in the paper is freely available online at https://github.com/BhallaLab/TimeCellAnalysis. The code is available as Extended Data 1.

## Acknowledgements

KGA and the experiments received support from Department of Biotechnology, BT/PR12255/MED/122/8/2016, to USB. NCBS-TIFR provided resources for analysis equipment and support to USB through intramural funds supported by the Department of Atomic Energy, Government of India, under project identification No. RTI 4006.

We thank Daniel Dombeck for help with the chronic preparation for physiological 2-P calcium recording of Hippocampal CA1 cells, and William Mau for his Temporal Information calculation code. We thank Vinu Varghese and Bhanu Priya Somashekar for helpful comments on the manuscript.

