## Supplementary material for "Synthetic Data Resource and Benchmarks for Time Cell Analysis and Detection Algorithms": All Supplementary Figures with Legends

Legends for Supplementary

Figure 1-1. Modulation profile along with the False Positive and False Negative rates per dataset, for important parameters configured in each of the 567 synthetic datasets generated. A-C: “Unphysiological Regime”, D-F: “Canonical Regime”, G-I: “Physiological Regime”.

Figure 6-1. A: Equivalence by XNOR matching the prediction lists from the top six detection algorithms (Blue: Time Cells; Red: Other Cells). B-F: Dependence of the predictive performance (F1 Score) on the various important synthetic dataset configuration parameters, B: Noise (%), C: Event Width (%ile), D: Imprecision (frames), E: Hit Trial Ratio (%), and F: Background Activity (𝜆).

Figure 6-2: Linear Regression fits for all algorithm parameter dependence curves with data points (red circles), best fit line (black), and the 95% prediction interval (PI; dotted black lines). The columns represent the physiology regime modulation parameter (out of the 5 main parameters tested), and the rows represent the various implemented algorithms for time cell detection.

Extended Data Figures (Supplementary)

Figure 1-1:

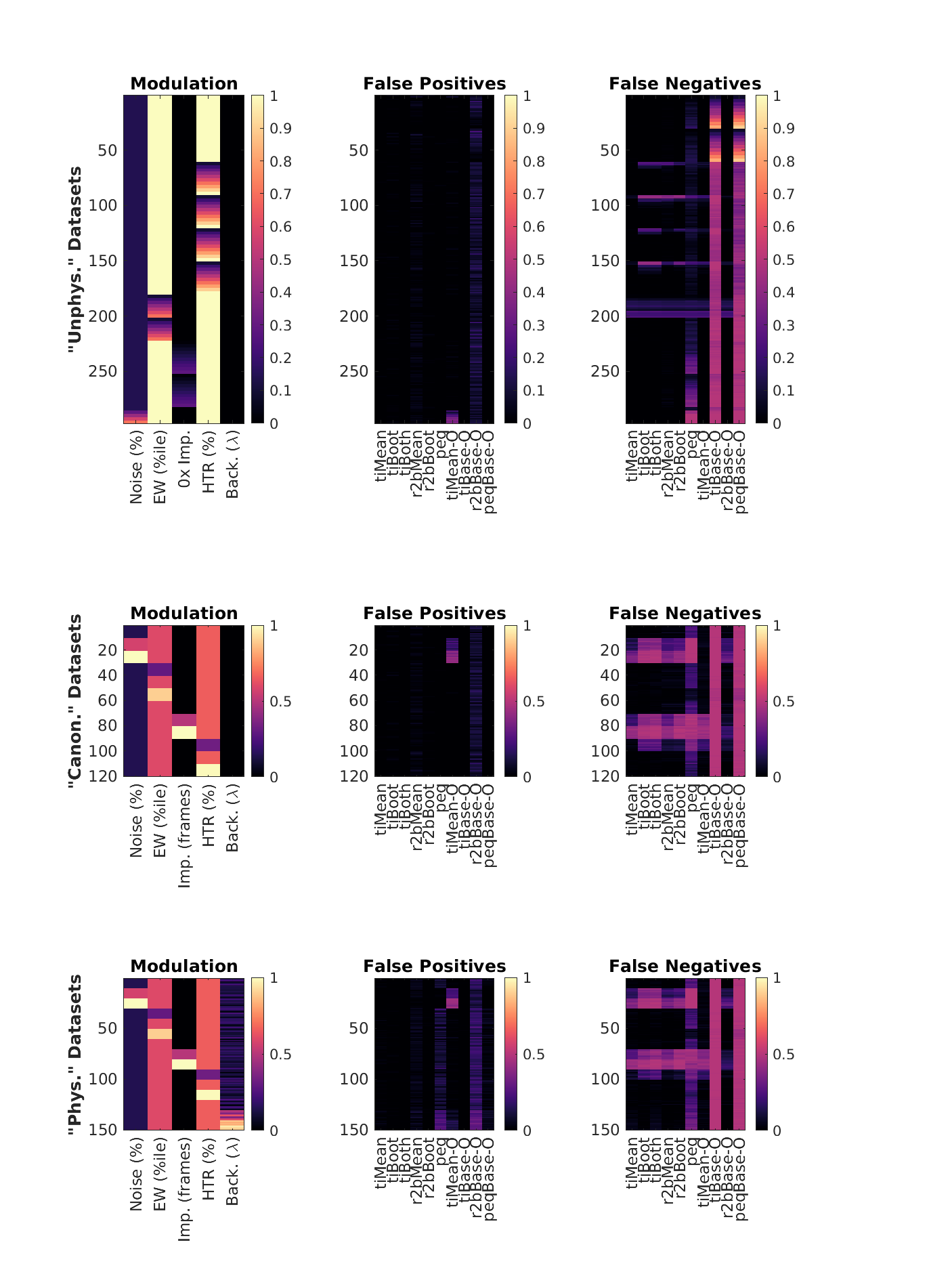

Figure 6-1:

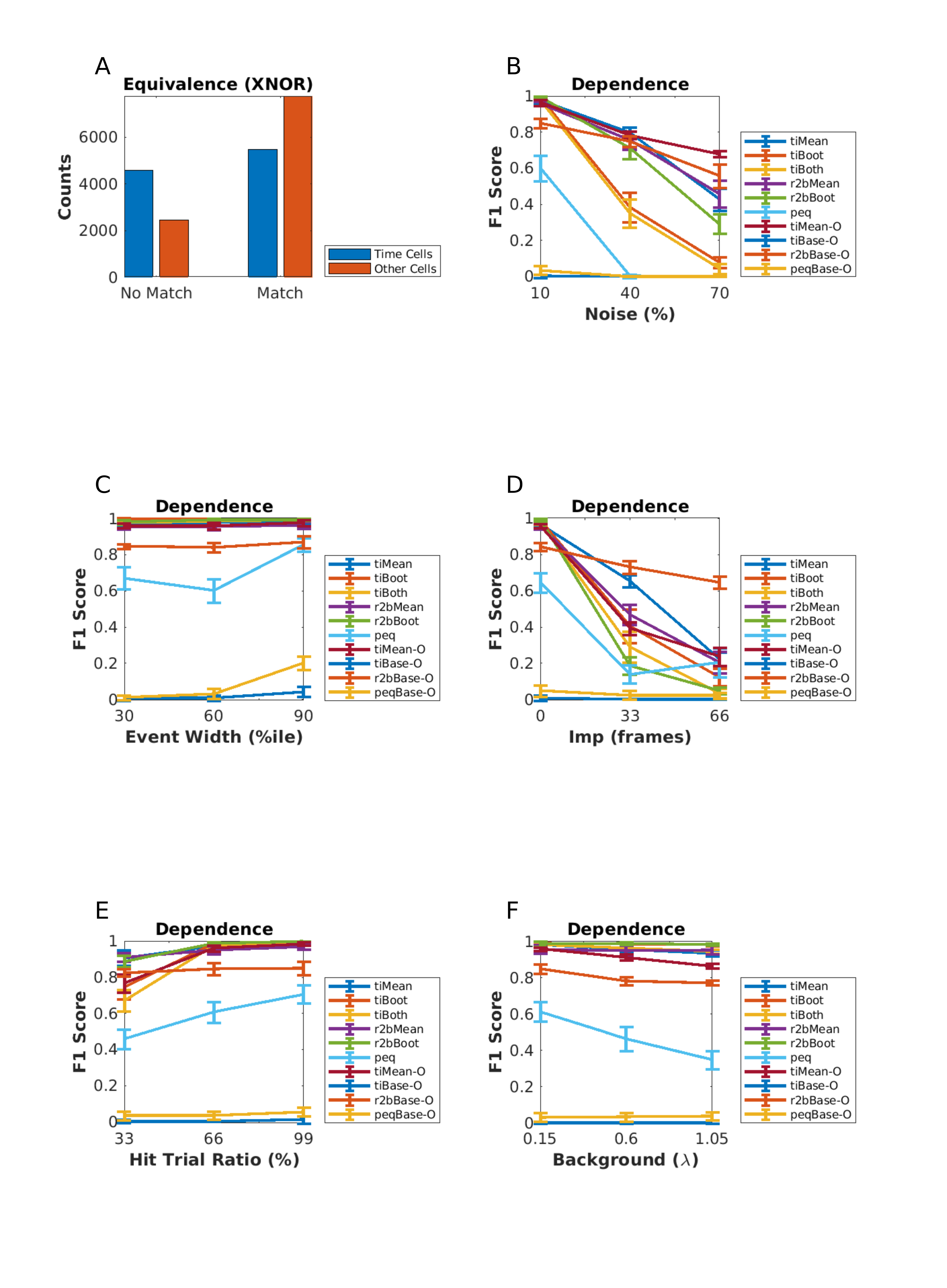

Figure 6-2:

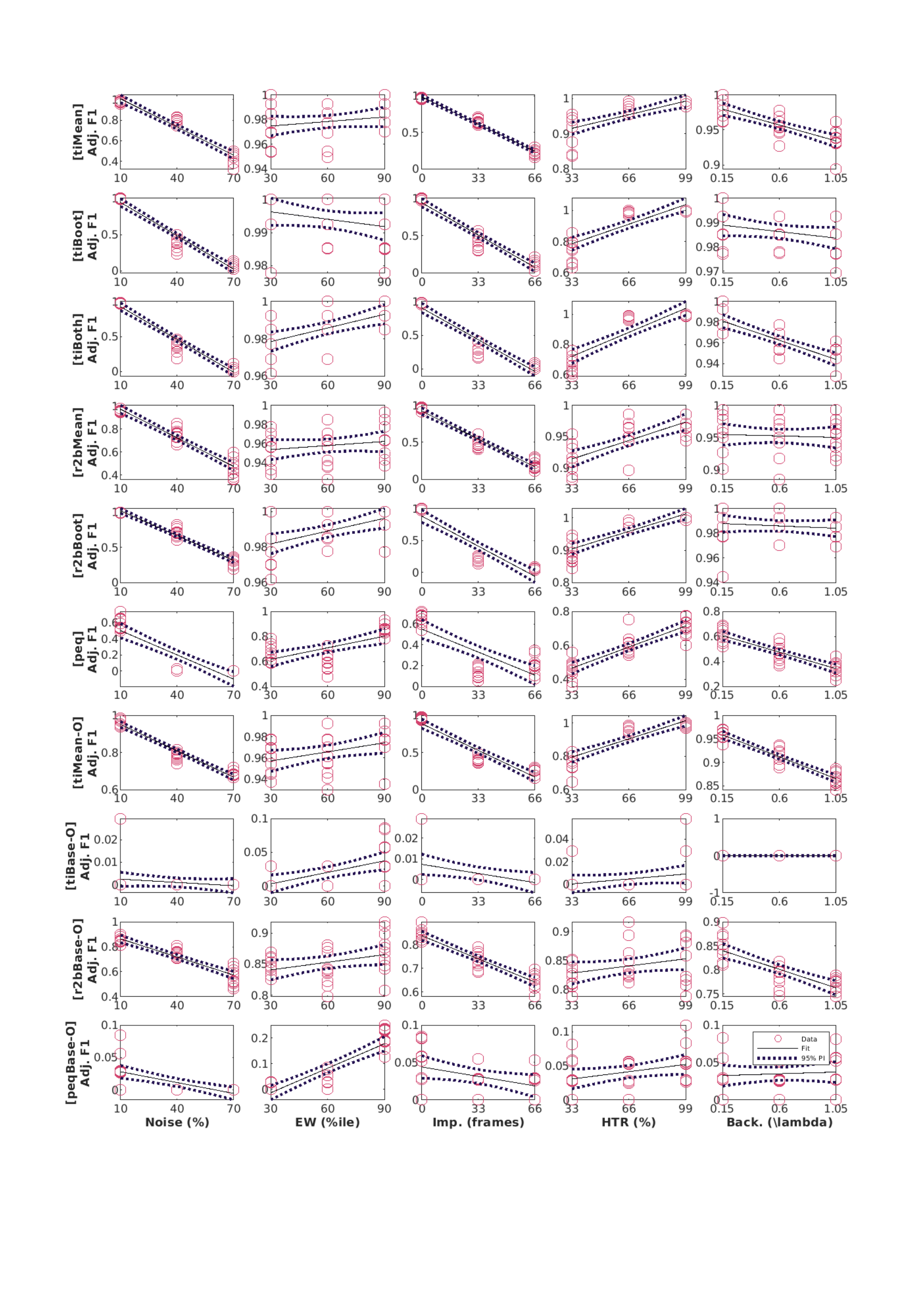
